# Left anterior temporal lobe is necessary for efficient lateralised processing of spoken word identity

**DOI:** 10.1101/510347

**Authors:** Thomas Cope, Yury Shtyrov, Lucy MacGregor, Rachel Holland, Friedemann Pulvermüller, James B Rowe, Karalyn Patterson

## Abstract

In the healthy human brain, the processing of spoken words is strongly left-lateralised, while the processing of complex non-linguistic sounds recruits brain regions bilaterally. Here we asked whether the left anterior temporal lobe, strongly implicated in semantic processing, is critical to this special treatment of linguistic stimuli. Nine patients with semantic dementia (SD) and fourteen age-matched controls underwent magnetoencephalography and structural MRI. Voxel based morphometry demonstrated the stereotypical pattern of SD: severe grey matter loss restricted to the left anterior temporal lobe. During magnetoencephalography, participants listened to word sets in which identity and meaning were ambiguous until utterance completion, for example *played* vs *plate*. Whereas left-hemispheric responses were similar across groups, patients demonstrated increased right hemisphere activity 174-294ms after stimulus disambiguation. Source reconstructions confirmed recruitment of right-sided analogues of language regions in SD: atrophy of left anterior temporal lobe was associated with increased activity in right temporal pole, middle temporal gyrus, inferior frontal gyrus and supramarginal gyrus. Moreover only healthy controls had differential responses to words *versus* non-words in right auditory cortex and planum temporale. Overall, the results indicate that anterior temporal lobe is necessary for normal and efficient processing of word identity in the rest of the language network.

## Introduction

The neural processing of spoken words is strongly lateralised to the dominant cerebral hemisphere, usually the left, while the processing of complex non-linguistic sounds recruits brain regions bilaterally (Zatorre *et al*, 1992, Shtyrov *et al*, 2000, Zatorre *et al*, 2002, Tervaniemi and Hugdahl, 2003). Across a range of primate species, acoustic information entering primary auditory cortex is rapidly transferred along reciprocal connections to the anterior temporal lobe (ATL) (Hackett, 2011, Friederici, 2012), a region that is strongly implicated in the representation and processing of semantic information in the human brain (Mummery *et al*, 2000, Pobric *et al*, 2007, Binney *et al*, 2010, Mion *et al*, 2010, Visser *et al*, 2010, Binder *et al*, 2011, Guo *et al*, 2013, Lambon Ralph *et al*, 2017). Here we asked whether the integrity of this region is necessary for the special, lateralised processing of spoken word identity.

This central question was motivated in part by a clinical observation. Neurodegeneration of the anterior temporal lobes, generally more severe in the dominant (usually left) hemisphere, results in the clinical syndrome of semantic dementia (SD, also known as the semantic variant of primary progressive aphasia type of frontotemporal dementia). SD erodes semantic memory and conceptual knowledge as well as language function (Warrington, 1975, Bozeat *et al*, 2000, Patterson *et al*, 2006), in keeping with emerging views of ATL as a transmodal semantic hub (Patterson *et al*, 2007, Guo *et al*, 2013, Lambon Ralph *et al*, 2017). In SD, processing of single spoken words is entirely adequate to enable repetition: if you ask an SD patient to repeat a long and complicated word like “hippopotamus”, they will typically do so correctly and effortlessly. But, ask the patient what a hippopotamus is, and the response from a mild case might be: “is it some sort of animal?” and from a moderate or severe case: “I don’t know”. Importantly, patients with SD may also struggle to repeat longer sequences of words or sentences, frequently displaying phonemic exchanges (e.g., “The *flag blew* in the wind” repeated as “The *blag flew* in the wind”), especially if the word sequence/sentence contains infrequently encountered words (Warrington and McCarthy, 1987, Patterson *et al*, 1994). Similarly, despite relatively preserved day-to-day episodic and prospective memory, patients with SD sometimes struggle on tests of delayed recall, producing answers that ‘sound-like’ the information they were asked to retain. A recent patient, asked to retain the name-and-address: “Harry Barnes, 73 Orchard Close, Kingsbridge, Devon”, recalled ten minutes later: “Harry Buns, 73 Awkward Close, I’ve forgotten the rest.”

These response patterns suggest that, with degeneration of the anterior temporal lobe, patients might be encoding information phonologically rather than lexically (Snowling *et al*, 1991, Gathercole, 1995). This leads to poorer recall performance for words that are no longer understood (Patterson *et al*, 1994, Knott *et al*, 1997), as patients lose the normal recall benefit for real words over non-words that is observed in healthy participants (Hulme *et al*, 1991). Indeed, there is evidence that in SD, the brain processing of real words and word-like nonwords becomes increasingly similar. For example, SD patients are impaired at distinguishing between real words and non-words in a visual lexical decision task, especially if the nonword in a word/non-word pair (such as FRUIT/FRUTE) follows a more typical orthographic pattern than the word, as measured by bigram and trigram frequencies (Rogers *et al*, 2004, Patterson *et al*, 2006). Similarly, patients with SD are relatively impaired at identifying acoustically degraded speech in categories for which they have impaired semantic knowledge (place names), compared to those for which their knowledge is in tact (number strings) (Hardy *et al*, 2018), and indeed generally show a striking advantage in verbal working memory for numbers compared to other word-types (Jefferies *et al*, 2004).

SD is characterised by progressive deterioration of conceptual knowledge, modulated by familiarity. Because it is a central semantic disorder, the cognitive impact is not confined to language; but language deficits are early and prominent, leading to an additional characterisation of the condition within the spectrum of primary progressive aphasias as semantic variant primary progressive aphasia (Gorno-Tempini *et al*, 2011). Deficits in confrontational naming and word comprehension are especially prominent, whereas repetition, grammar, and motor speech are usually well preserved until late in the illness. The syndrome results from neurodegeneration of anterior temporal lobes that is usually more severe in the left hemisphere, and is almost always caused by TDP-43 type-C neuropathology (Hodges *et al*, 2009, Rohrer *et al*, 2011, Spinelli *et al*, 2017). By the time of clinical presentation, this localised neurodegeneration is usually already severe, and even patients at a moderate stage of illness and living relatively normal daily lives may show 50-80% loss of left anterior temporal grey matter (Hodges and Patterson, 2007).

The fact that SD patients can perform an ‘off-line’ task like listening to and repeating a spoken word does not establish that the earliest stages of spoken-word processing in SD are unaltered. In the healthy brain, early processing, whilst not unilateral, is biased towards the left hemisphere with increasing left-lateralisation observed as information moves forward from posterior to anterior regions (Marinkovic *et al*, 2003). Here we recorded neural activity with magnetoencephalography (MEG) while participants listened to word sets in which word identity and meaning were ambiguous until utterance completion.

We compared neural responses between healthy participants and people with neurodegeneration of ATL from SD. The advantage of MEG in this context is that it allowed us to compare the time-course of neural activity between these two groups with sufficient spatial resolution to assess the approximate location of several simultaneously-active brain regions. MEG has been shown to be sensitive to both semantic decisions (Hughes *et al*, 2011) and auditory change detection abnormalities (Hughes *et al*, 2013, Hughes and Rowe, 2013) in frontotemporal dementia. Here we employed a spoken-word version of the auditory mismatch paradigm (for a review see Näätänen *et al*, 2007), in which repeated ‘standard’ words (for example PLAY) were changed in either tense or semantic meaning by the spliced addition of the additional ending syllables d/t (to become, in this case, PLAYED or PLATE). This paradigm is a sensitive tool for measuring automatic lexico-semantic processing of spoken words in the brain (Pulvermüller *et al*, 2006, Shtyrov *et al*, 2010) and has a special benefit for patient studies as it does not require any active stimulus processing, or even attention on the auditory stream (Gansonre *et al*, 2018). Here, presentation was designed such that the occurrence and timing of a deviant word were predictable, but the identity and meaning of the word were unpredictable until the last tens of milliseconds of its utterance. The processing of deviant word-endings in this paradigm recruits language-specific brain regions, and has previously been demonstrated to produce a strongly left-lateralised response in young, healthy listeners (Holland *et al*, 2012). This left-lateralisation is most prominent in the first few hundred milliseconds, time windows that are generally considered to reflect the auditory analysis of linguistic information (MacGregor *et al*, 2012). Later cognitive processing of the meaning of language, first reflected in the N400 (300-500ms) response, is typically symmetric over the hemispheres or even right lateralized (Kutas and Federmeier, 2011).

We tested directly how the pattern of neural activity involved in processing spoken words is affected by disruption of the reciprocal connectivity between undamaged early auditory regions in posterior superior temporal lobe and severely compromised transmodal semantic regions in ATL. Specifically, our analyses of the MEG data from SD patients relative to healthy age-matched controls addressed three questions: whether degeneration of the left ATL would result in:

1. disruption of the normal pattern of laterality in spoken word processing, in particular, a shift from a left-dominant pattern in controls to more bilateral activations?
2. increased similarity in the distribution of the brain response to words and word-like non-words?
3. an overall shift in brain activity from areas implicated in word processing to those involved in the analysis of non-linguistic acoustic features?

## Methods

### Ethics

All study procedures were approved by the UK National Research Ethics Service. All participants had mental capacity and gave written informed consent to participation in the study.

### Participants

Eleven patients with semantic dementia (SD) were recruited from a single tertiary referral cognitive clinic. All patients met consensus diagnostic criteria for both SD (Neary *et al*, 1998) and semantic variant primary progressive aphasia (Gorno-Tempini *et al*, 2011). Nine of the patients (eight right-handed, one left-handed) tolerated the MEG environment sufficiently to complete the whole experimental paradigm, and provided the data reported here. Eight were able to undertake a research structural MRI brain scan.

Fourteen right-handed, healthy individuals of a similar age were recruited as controls. All produced complete MEG datasets and underwent a structural MRI head scan.

Participant demographics are shown in Table 1.

**Table 1:**
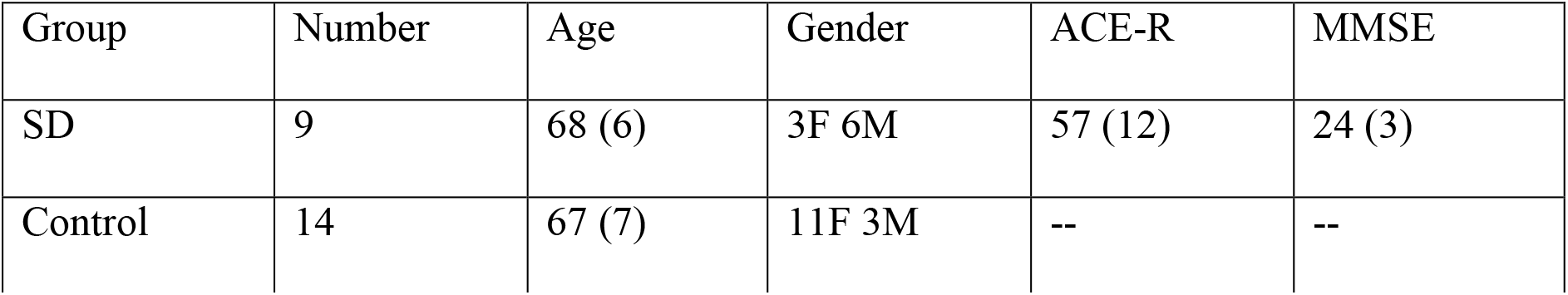
Participant demographics. Mean (standard deviation). ACE-R = Addenbrooke’s cognitive examination, revised edition. MMSE = Mini-Mental State Examination

### Experimental paradigm

The procedure closely mirrored that of a previously published MEG study of the hemispheric laterality of word processing in healthy young adults (Holland *et al*, 2012). Participants sat upright in a magnetically shielded room, watching a silent movie while passively listening to spoken words delivered through an in-ear air tube system. No response was required, thereby reducing the difficulties inherent in the explanation and retention of a behavioural task for patients with semantic impairment.

Words consisted of one of three standard (template) words and two deviants for each standard that varied in their endings (*Figure 1*). Standards comprised the real words ‘PLAY’ and ‘TRAY’, and the pseudo-word ‘KWAY’, all closely matched acoustically and phonetically. Deviant endings were created by the spliced addition of /d/ or /t/ to the end of a standard word, avoiding co-articulation effects and resulting in the six deviant stimuli ‘PLAYED’, ‘PLATE’; ‘TRADE’, ‘TRAIT’; and ‘KWAYED’ (or ‘KWADE’), ‘KWATE’. This acoustic splicing avoided co-articulation effects, and resulted in a divergence point between /d/ and /t/ endings 10ms after the offset of the standard word.

**Figure 1:**
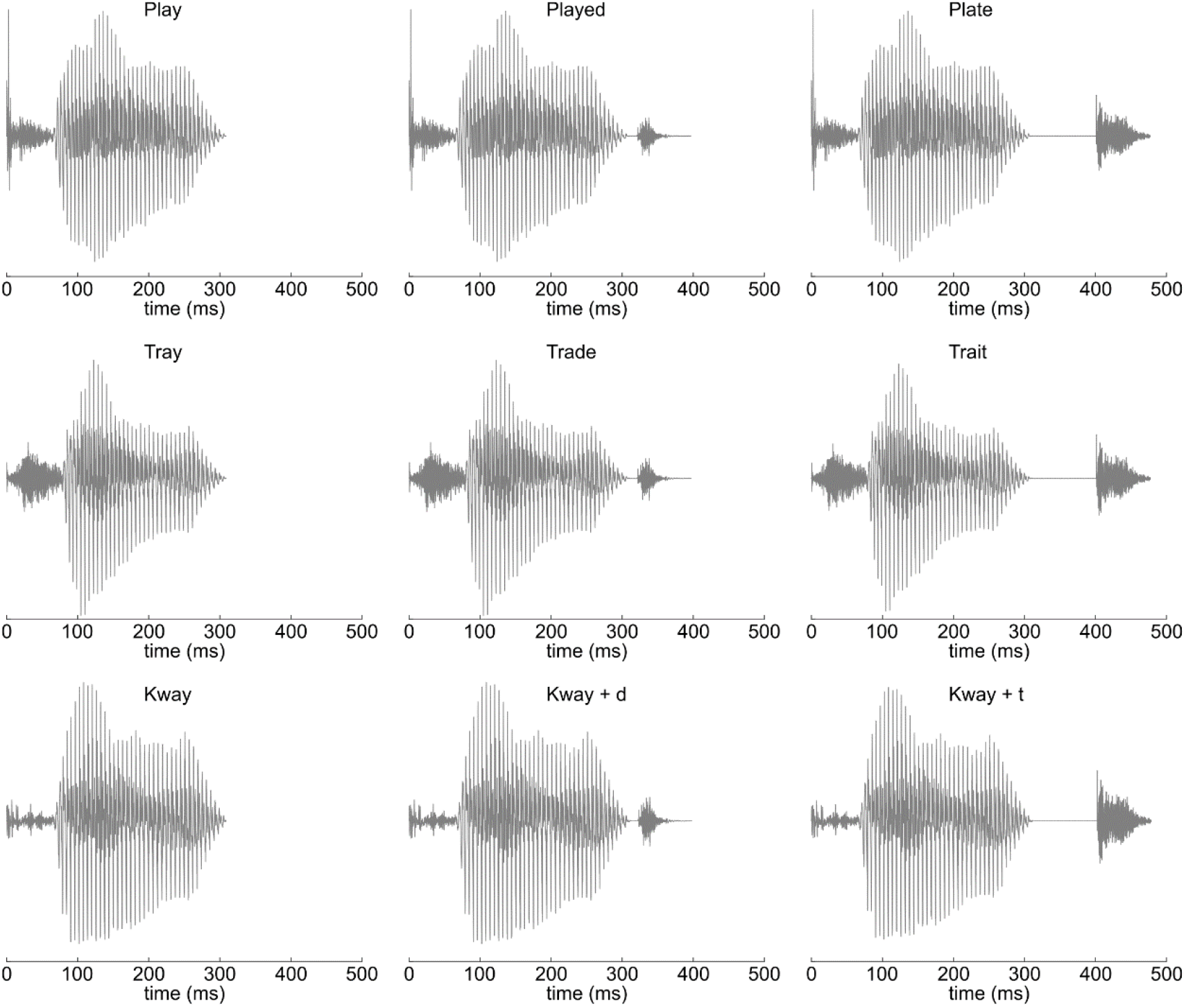
Waveforms of the three standard words, with the spliced addition of /d/ and /t/ deviant endings. All stimuli within each triplet were identical for the first 320ms.

Presentation followed a repeating pattern of 4 standards to 1 randomly chosen deviant, with a fixed 1 second inter-onset-interval, such that the *occurrence* of a deviant was entirely predictable but its *identity* was not. For example, after four presentations of the word ‘PLAY’, the next word would be either ‘PLAYED’ or ‘PLATE’. Stimuli were presented in blocks such that each participant heard a single template word 800 times and each of its deviant forms 100 times. Blocks therefore lasted 1000 seconds (approximately 17 minutes), and the order of presentation was counterbalanced across participants.

### Voxel Based Morphometry

Eight patients with SD and 14 controls underwent structural MR imaging using a 3T Siemens Magnetom Tim Trio scanner with a 32-channel phased-array head coil. A T1-weighted magnetization-prepared rapid gradient-echo (MPRAGE) image was acquired with repetition time (TR)=2250-2300ms, echo time (TE)=2.86-2.98ms, in-plane resolution of 1.25×1.25mm, 1.25mm slice thickness, inversion time=900ms and flip angle=9°.

Voxel based morphometry analysis used SPM12 (www.fil.ion.ucl.ac.uk/spm). Images were first approximately aligned by coregistration to an average image in MNI space, before segmentation and calculation of total intracranial volume (TIV). After segmentation, a study-specific DARTEL template was created from the 8 patient scans and the 8 controls mostly closely matched in age on a patient by patient basis, using default parameters. The remaining controls were then warped to this template. The templates were affine aligned to the SPM standard space using ‘Normalise to MNI space’ and the transformation applied to all individual grey-matter segments together with an 8mm FWHM Gaussian smoothing kernel. The resulting images were entered into a full factorial general linear model with a single factor having two levels, and age and TIV as covariates of no interest. This model was estimated in the classical manner, based on restricted maximum likelihood. Voxels were defined as atrophic if they were statistically significant at the cluster FWE p<0.05 level, with an uncorrected cluster defining height of p<0.001.

### Magnetoencephalography data acquisition and pre-processing

MEG data were acquired with a 306-channel Elekta Neuromag Vectorview system with 102 magnetometers and 204 paired planar gradiometers. Data were digitally sampled at 1kHz and high-pass filtered above 0.01Hz. Throughout scanning, the 3D position of five evenly distributed head position indicator (HPI) coils was continuously monitored relative to the MEG sensors. The positions of these indicator coils, relative to overall head shape and the position of three anatomical fiducial points (nasion, left and right pre-auricular), were measured before scanning with a 3D digitiser (Fastrak Polhemus). Electrooculography data were also acquired to allow later data artefact removal.

MEG and HPI data were pre-processed in Neuromag Maxfilter 2.2 to perform Signal Source Separation (Taulu *et al*, 2005) for motion compensation and environmental noise suppression. All subsequent data analysis steps were undertaken in Matlab 2013a (The Mathworks Inc., 2015) using the software packages SPM12-r6906 (Wellcome Trust Centre for Neuroimaging, London, UK), FieldTrip (Donders Institute for Brain, Cognition, and Behavior, Radboud University, Nijmegen, The Netherlands) and EEG lab (Swartz Center for Computational Neuroscience, University of California San Diego). Magnetometer and planar gradiometer data were subjected to separate indepenxg121 cl:1477dent component analyses for artefact rejection. Artefactual components were automatically identified by a conjunction of temporal correlation with electrooculography data and spatial correlation with separately acquired template data for blinks and eye movements.

The cleaned data were then sequentially epoched from-500 to 1500ms relative to word onset; downsampled to 250Hz; baseline corrected to the 100ms before word onset; lowpass filtered below 40Hz; merged across recording session; averaged using the SPM robust averaging algorithm, which produces an average after weighting individual epochs according to their consensus; and re-filtered below 40Hz to remove high frequency components introduced by robust averaging. Planar gradiometer data pairs were root-mean-square combined; converted to scalp-time images; smoothed with a 10mm spatial kernel and 25ms temporal kernel; and finally masked for statistical analysis to time windows from -100ms to 900ms relative to word onset and -100ms to 600ms relative to the timing of standard word offset.

### Sensor-space evoked analysis

The initial analysis of the contrast between standard and deviant words was undertaken in sensor space, for which the signal to noise ratio is higher than data in source space (Martín - Buro *et al*, 2016) and no a-priori specification of time windows of interest is required. To allow robust interpretation of laterality effects, this analysis was performed on the planar gradiometer data, for which signal magnitude at the scalp is maximal directly over the source of neural activity (Parkkonen, 2010). A flexible factorial design was specified in SPM12, and interrogated across all participants for main effects of interest. The scalp location of peak statistical effect was identified on each side (left and right; in all cases p(FWE) was < 0. 01). The time-courses of the sensor data extracted at each of these scalp locations was then compared across groups at every time point. This approach is superior to the extraction of time-courses from a single, closest gradiometer pair, as it inherently controls for interindividual differences in head position relative to the detector array, and by virtue of spatial smoothing includes weighted information from nearby sensors, reducing the effect of differential noise in any one superconducting quantum interference device (SQUID). It does not represent double dipping as the location of interest for between-group comparison was defined by the orthogonal contrast of overall main effect (Friston and Henson, 2006, Kriegeskorte *et al*, 2009, Kilner, 2013). When comparing extracted time-courses, a significant group × condition interaction was defined as at least seven consecutive timepoints of p < 0.05, resulting in a sustained effect over >=28ms, exceeding the temporal smoothing induced by lowpass filtering at 40 Hz.

Laterality effects in the analysis of deviant word endings were assessed through laterality quotients (Holland *et al*, 2012). These were calculated for every time-point outside of the baseline period for each individual separately as:

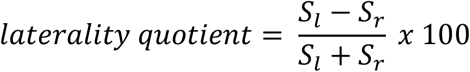

Where *S_l_* and *S_r_* are the magnitudes of the deviance effect at the same scalp locations as interrogated for the group by deviance interaction on each side. The laterality quotients were assessed at every time point for both difference from zero for each group separately, and for group by deviance interactions.

### Source-space evoked analysis

Source reconstructions were undertaken in SPM12 for two purposes. The first was the intention to localise the brain basis of any neurophysiological interaction between word ending and group that was statistically demonstrated in sensor space. The second was to enable primary source-space statistical analysis of standard-word-identity effects and interactions (where four times as many repetitions of each stimulus and a precisely-defined temporal window of interest based on overall contrast magnitude allowed adequate signal-to-noise ratio for a primary source-space analysis to be undertaken).

Single shell MEG forward models were created for each participant based on individually recorded head shapes co-registered to MRI scans using fiducial points. Magnetometer and planar gradiometer data were combined (Henson *et al*, 2009) and group source inversion across all participants was undertaken with sLORETA (Pascual-Marqui, 2002) across epochs of -100ms to 900ms relative to spoken word onset. Within the time window of interest, condition estimates were computed in a 1-40Hz frequency band and converted into images. These images were then subjected to statistical analysis within a flexible factorial general linear model design identical to that employed for the sensor-space evoked analysis. This led to the creation of t-score maps contrasting the neural response to standard and deviant words, which were then thresholded for visualisation.

## Results

### Voxel Based Morphometry

Voxel based morphometry (*Figure 2*) demonstrated the expected pattern of SD, with predominant grey matter loss compared to the control group in the left ATL (peak [-29 1 -40] t(18)=13.34 FWE p<0.001), with atrophy of the same region on the right that was less marked in magnitude and extent (peak [36 14 -32] t(18)=8.05 FWE p=0.004). Volume loss of the left insula was also observed that exceeded the cluster defining height (as illustrated in Figure 2) but was not significant at the corrected voxel level (peak [-33 14 8] t(18)=5.12 FWE p=0.29). Grey matter volume elsewhere was not statistically different from control participants.

**Figure 2:**
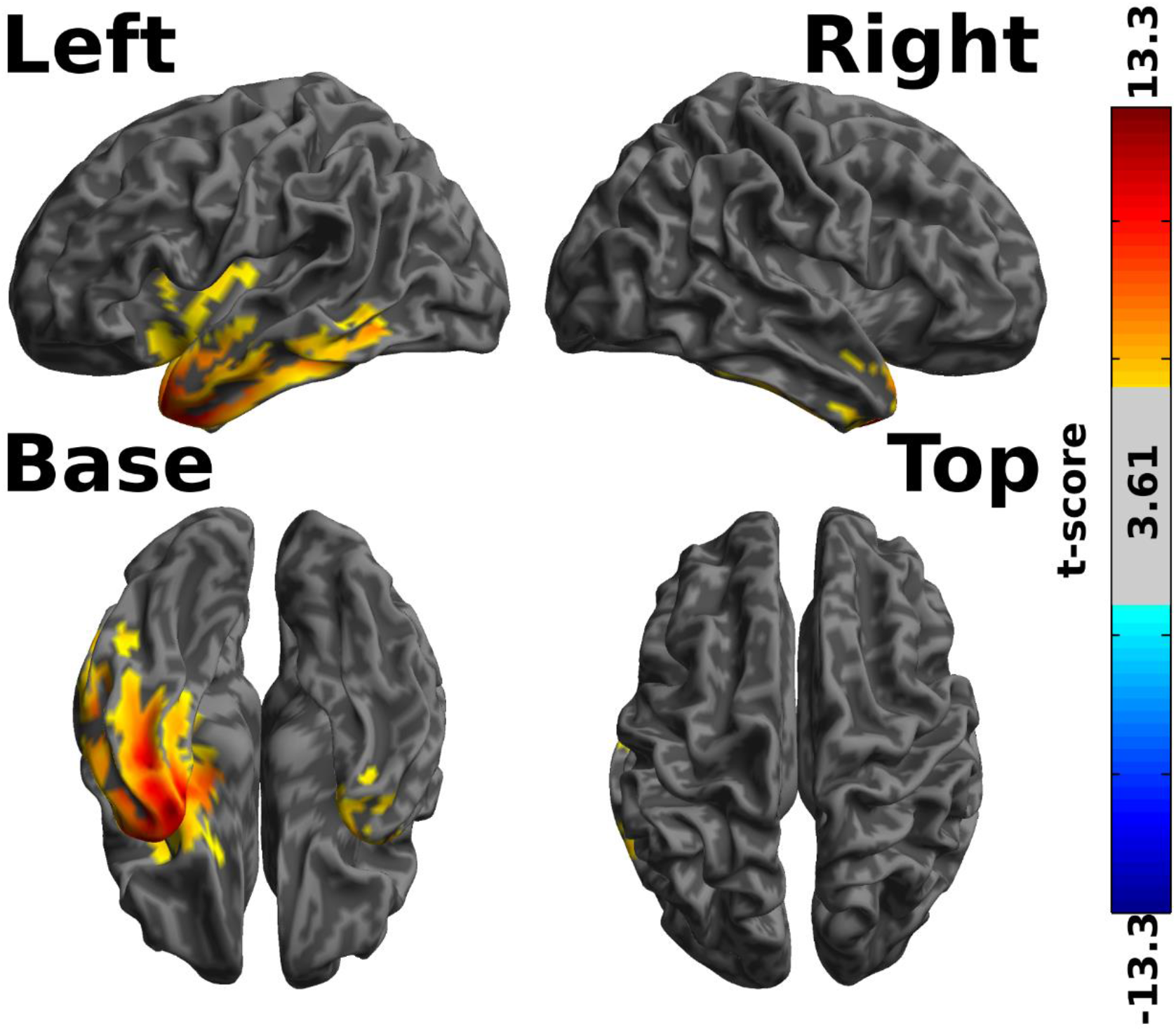
Voxel based morphometry statistical comparison of 8 participants with SD against 14 age-matched controls. Shaded areas represent t-scores for greater grey matter volume in the control group, cluster thresholded at FWE p<0.05 with a height threshold at uncorrected p<0.001. No voxels demonstrated greater grey matter volume in the patient group. Statistical maps are overlaid onto the same partially inflated template brain used to illustrate the MEG source reconstructions in figures 5 and 7.

### Overall magnetic response to standard words

At the scalp locations of peak response overlying each hemisphere ([-42 -9] on the left, [42 - 9] on the right, roughly overlying superior temporal lobe on each side), overall magnetic response to the three standard stimuli (2 words and 1 non-word) was significantly greater in the control group than the SD group in an early (36-72 ms) and a late (112-352 ms) time window relative to word onset (*Figure 3 upper*). The distribution of this response was similar across the two groups (*Figure 3 lower*).

**Figure 3:**
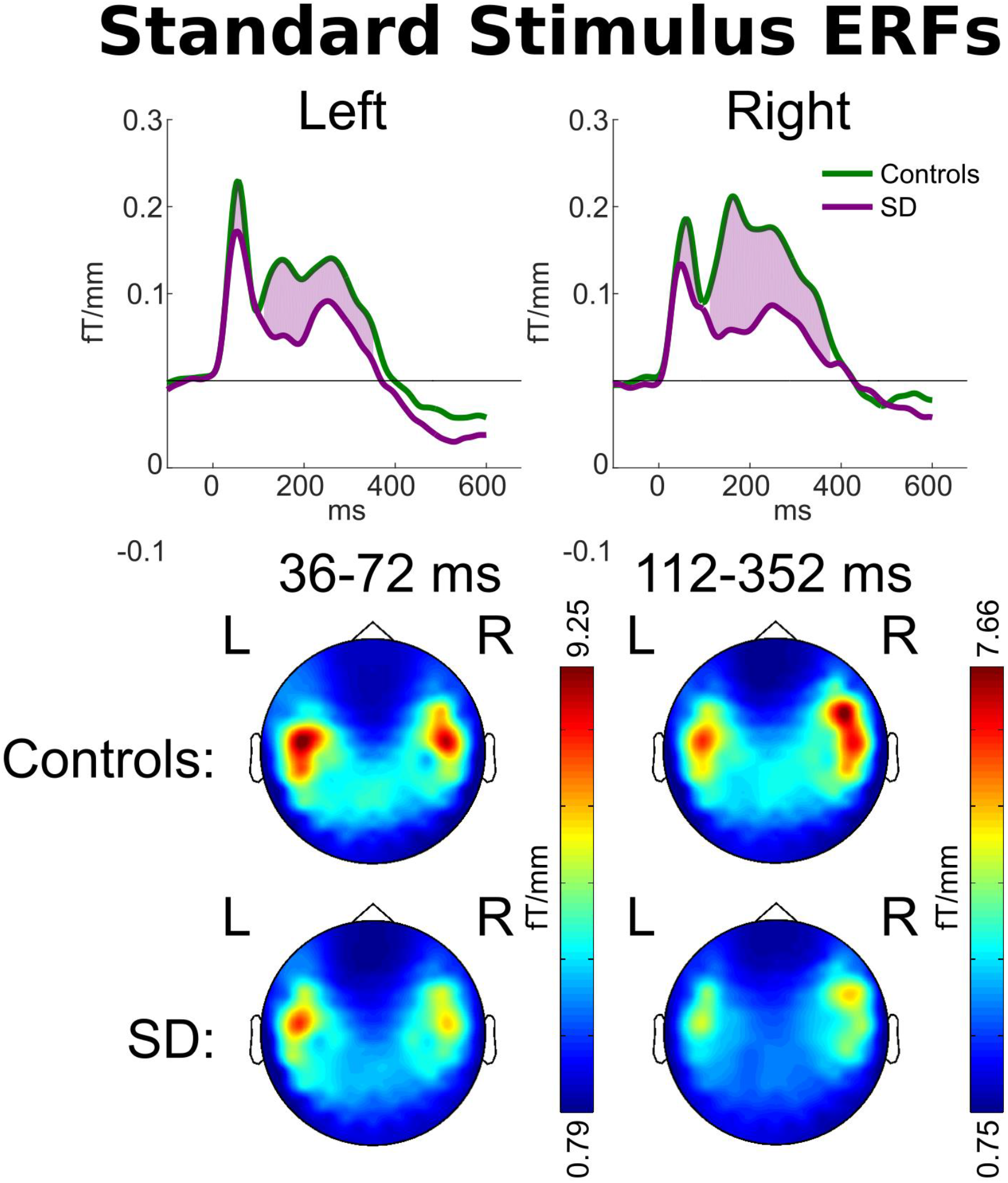
Upper: Magnetic field power recorded by planar gradiometers at the scalp locations of peak overall response to the standard word overlying each hemisphere. Responses are time-locked to word onset. Purple shading indicates time periods at which a statistical difference was observed in signal magnitude between patients and controls. Lower: Scalp signal topographies for each group, averaged within each period of statistically significant difference.

### Response to deviant word disambiguation

Despite the group difference in overall magnetic power in response to standard words, both groups demonstrated peak responses to the overall contrast between standard and deviant word endings of similar magnitude, with a much larger response to deviant words at around 100-160ms after stimulus disambiguation (*Figure 4 upper*). As has been previously observed in younger participants (Holland *et al*, 2012), for the older controls the deviance response for word ending was significantly greater on the left than on the right during this early peak. Indeed, for controls the laterality quotient was significantly greater than zero (more activity on the left) for every time point from 128-440ms (t(13) p<0.05; peak t(13) = 7.71, p=3.36×10^−6^ at 256ms). While patients demonstrated a deviance response of very similar average magnitude during this early time window (lines almost overlapping on both sides before 150ms in *Figure 4 upper*), due to the smaller group size and greater between-individual variability, the patient laterality quotient did not significantly differ from zero at any time point.

At later time windows (184-304ms after standard word offset, which is 174-294ms after the divergence point between /d/ and /t/ endings), a significant group by deviance interaction was observed in the right hemisphere, such that patients with SD demonstrated a larger difference between deviant and standard stimuli in the right hemisphere (peak t(21)=3.13, p=0.0050). Scalp topographies of average power during this period (*Figure 4 lower*) confirmed that this was not an effect restricted to the peak location, but rather represented a more general shift from highly left lateralised responses in controls to bilateral processing in patients with SD. Indeed, patients and controls demonstrated significantly different laterality quotients between 232-292ms after standard word offset (peak t(21)=2.90, p=0.0086 at 256ms).

**Figure 4:**
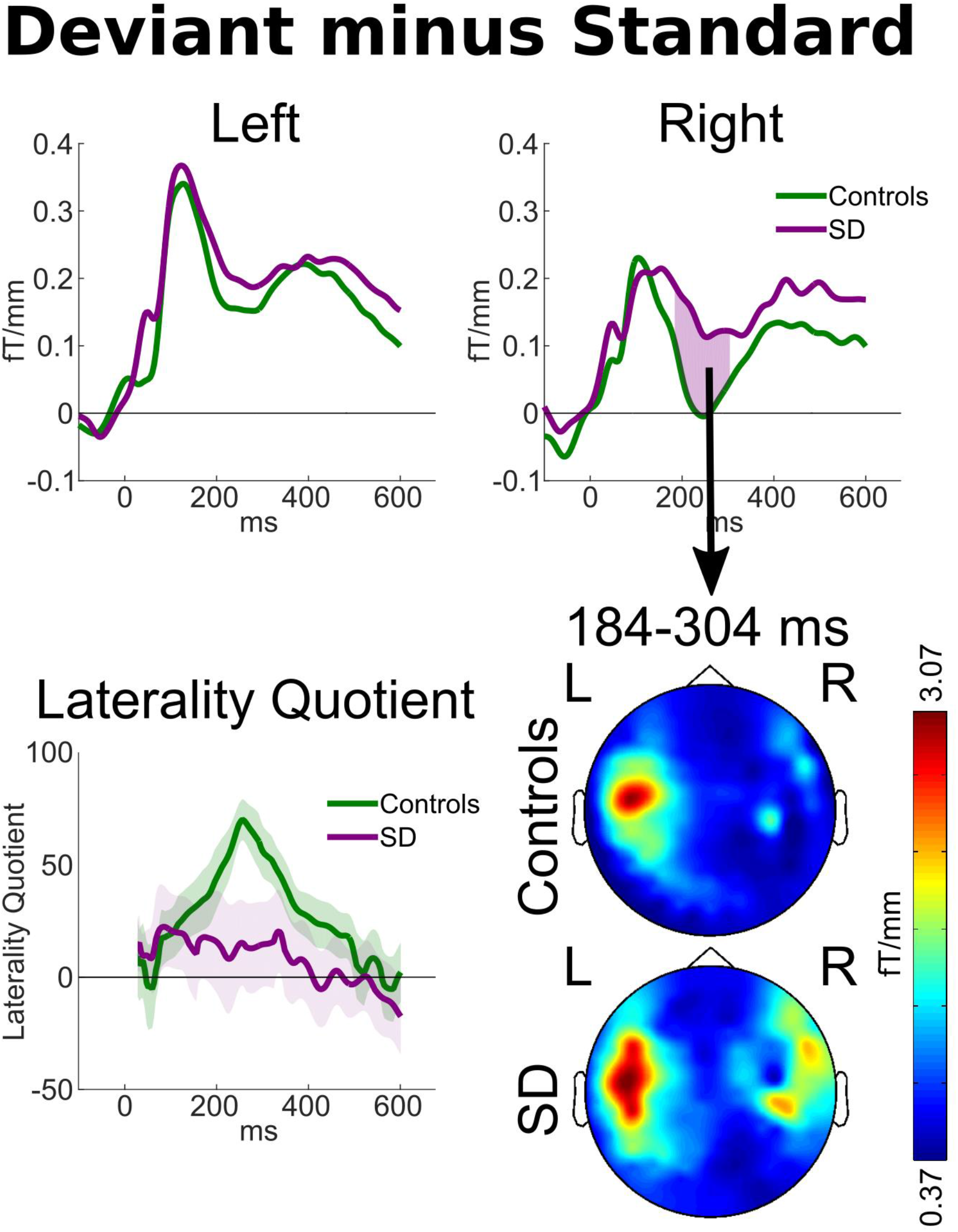
**Upper:** Magnetic field power recorded by planar gradiometers at the scalp locations of overall peak contrast between the average responses to all deviant words minus all standard words, relative to standard word offset. Pink areas indicate statistically significant group by deviance interactions as defined by p<0.05 sustained for ≥ 7 samples, exceeding the duration of temporal smoothing. **Lower left:** The laterality quotient of the deviance response for each group. Calculated such that fully left sided deviant responses would be +100, fully right sided responses -100. The shaded areas around each line encompass +/-one standard error. **Lower right:** Scalp signal topographies for each group, averaged across the period of the statistically significant group by condition interaction observed in the peak right-sided sensor.

We performed source localisations to assess the brain basis of the group difference in deviance response that we have statistically demonstrated in sensor space. Consistent with the scalp topographies in *Figure 4*, between 240-280ms after standard word offset healthy controls demonstrated a highly lateralised response predominantly involving left planum temporale and parietal lobe, with some involvement of inferior frontal regions (*Figure 5 upper*). Patients with SD demonstrated similar left sided responses, which were of lower average magnitude than controls, but not to a statistically significant degree. However, they had much more extensive activation of the right hemisphere (*Figure 5 middle*), again consistent with the sensor-space results presented in *Figure 4*. The voxelwise group by condition contrast (*Figure 5 lower*) demonstrated above-threshold clusters, with peak differences assessed by the Neuromorphometrics atlas to be in right temporal pole ([48 14 -2] t(357)=4.62), right middle temporal gyrus ([48 -32 -6] t(357)=4.15), right frontal operculum ([54 16 26] t(357)=4.04), right inferior temporal gyrus ([54 -44 -26] t(357)=3.50), and right supramarginal gyrus ([56 -28 46] t(357)=3.21), in what might be deemed right sided analogues of a classical map of the brain regions involved in language (Friederici *et al*, 2017). In all cases where these right-hemispheric differences were observed, patients with SD demonstrated equal or greater modulation of brain activity as a function of word ending than controls, despite the lower overall power of their magnetoencephalography response to spoken words (*Figure 3*).

**Figure 5:**
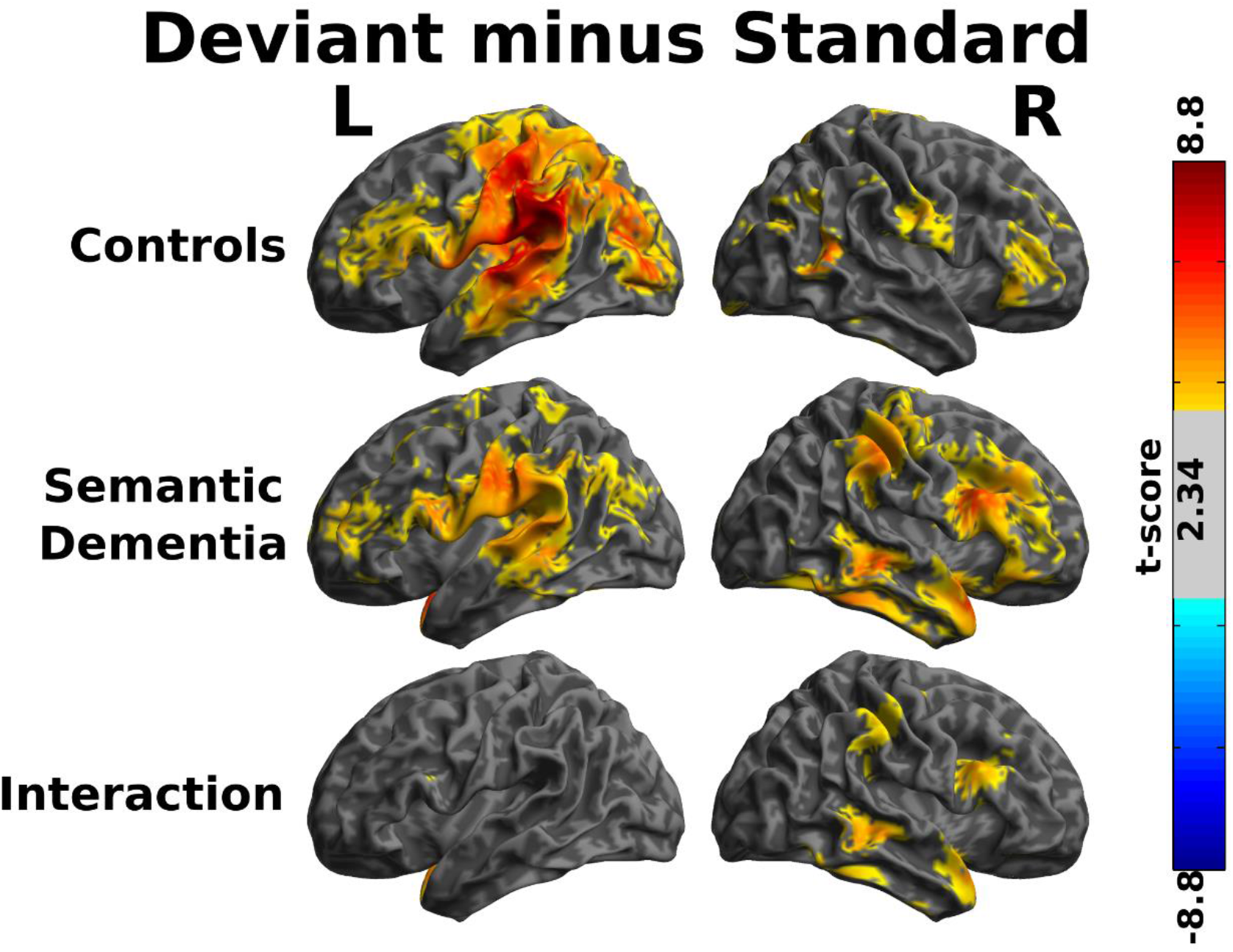
Source reconstructions of the contrast between standard and deviant words between 240-280ms after the offset of the standard word, the time window during which the largest group by deviance interaction was demonstrated in sensor space (cf Figure 4). Shaded areas represent t-scores thresholded for visualisation at t>2.34 (equivalent to uncorrected p<0.01). Two-tailed statistical tests were performed, but all surviving contrasts were greater in the deviant than the standard, and (for the third panel) the effect of deviance was greater in the patients than the controls.

### Response differences according to standard word identity

We explored the consequences of left ATL neurodegeneration for the neuronal processing of standard words. An initial assessment of magnetic response power across all gradiometers revealed that the only large differences between word pairs occurred in early time windows, between 50 and 70ms after word onset (*Figure 6*), corresponding to the early peak in overall brain response (*Figure 3 upper*).

**Figure 6:**
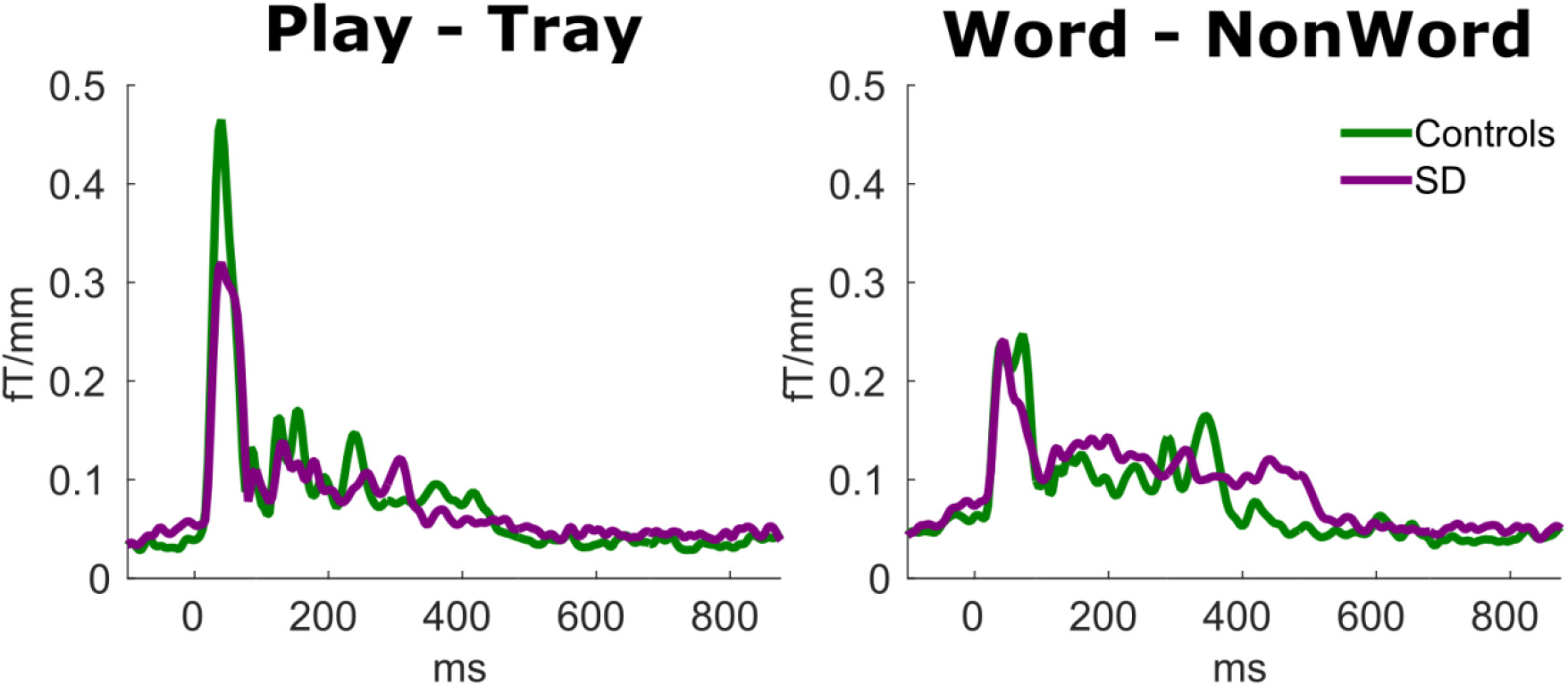
Overall difference in magnetic power detected by planar gradiometers across the whole scalp as a function of word identity, relative to the onset of the standard word.

To test for the presence of interactions between group and standard word identity we therefore performed source reconstructions in the 50-70ms time window and interrogated these with a primary whole-brain SPM.

For the contrast between real words (PLAY + TRAY) and the non-word KWAY (*Figure 7*), controls displayed significantly greater brain activity for non-words than words bilaterally (*Table 2*). The only significant interaction between this contrast and group was in right hemisphere regions surrounding primary auditory cortex and planum temporale in superior temporal lobe. Controls demonstrated a significantly greater difference in activity between words and non-words in these regions than patients with SD. Indeed, although the time window of reconstruction was chosen to capture the peak neuronal response across the whole brain, no individual voxels survived statistical thresholding in the SD group. In this group the non-significant locations of peak contrast in each hemisphere were: left postcentral gyrus at MNI [-60, -6, 22], t(357)=3.56, p(FWE)=0.43, and right frontal operculum at MNI [46, 14, 8], t(357)=3.92, p(FWE)=0.16.

For the contrast between the verb PLAY and the noun TRAY, participant group by word identity interactions were demonstrated in both directions (*Figure 8*, *Table 2*). All interactions were in the left hemisphere. Patients with SD displayed a greater effect of word identity in regions surrounding primary auditory cortex and planum temporale in posterior superior temporal lobe. Controls, on the other hand, demonstrated a greater effect of word identity around supramarginal gyrus in parietal lobe.

Overall, therefore, both groups were characterised by a left-lateralised effect of real word identity, but group by word interactions revealed that in SD this was greater around primary auditory regions in temporal lobe while in controls it was greater around parietal regions that some researchers have proposed to play a significant role in linking phonological analysis to meaning (Robson *et al*, 2013, Robson *et al*, 2017). Further, relative to the SD group, controls displayed a significantly greater response to non-words than to real words across superior temporal lobe and planum temporale, especially on the right. Indeed, source reconstructions demonstrated no significant clusters of activity in the SD group that differed between real words and non-words.

**Figure 7:**
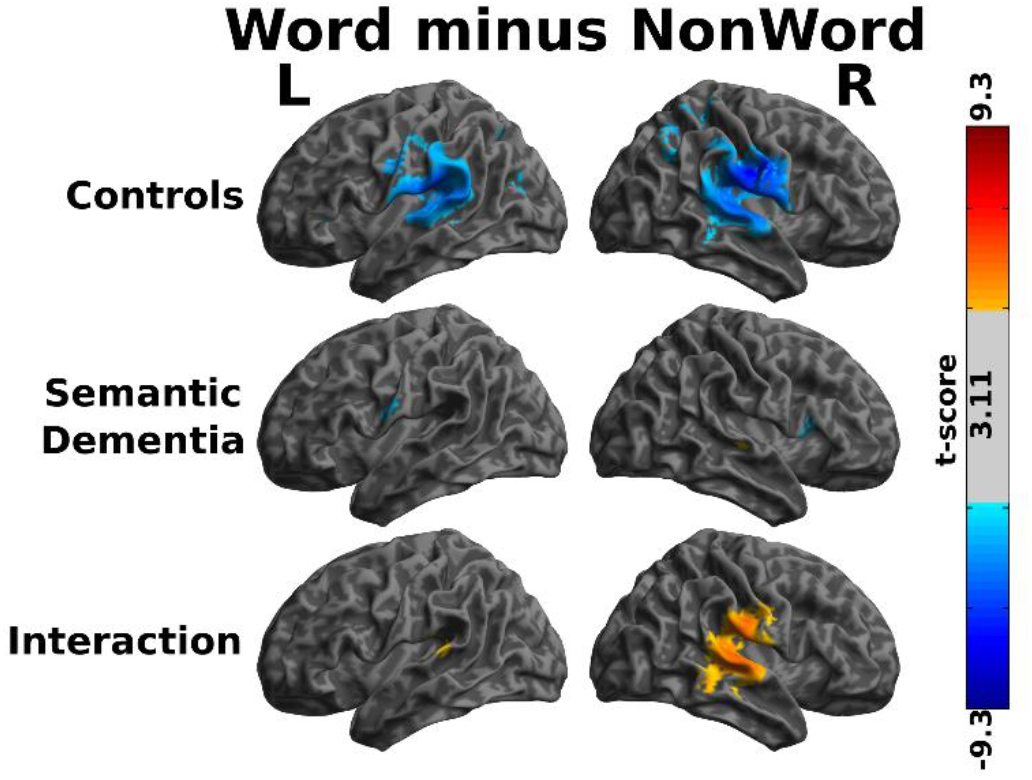
Source reconstructions of the contrast between standard real words and non-words 50-70ms after onset, the time window during which the largest effect of word identity was demonstrated across the whole brain (cf Figure 6). Shaded areas represent t-scores thresholded at uncorrected p<0.001 (t>3.11). In the lower right panel red shaded regions represent greater contrast in controls; no voxels demonstrated greater contrast in SD.

**Figure 8:**
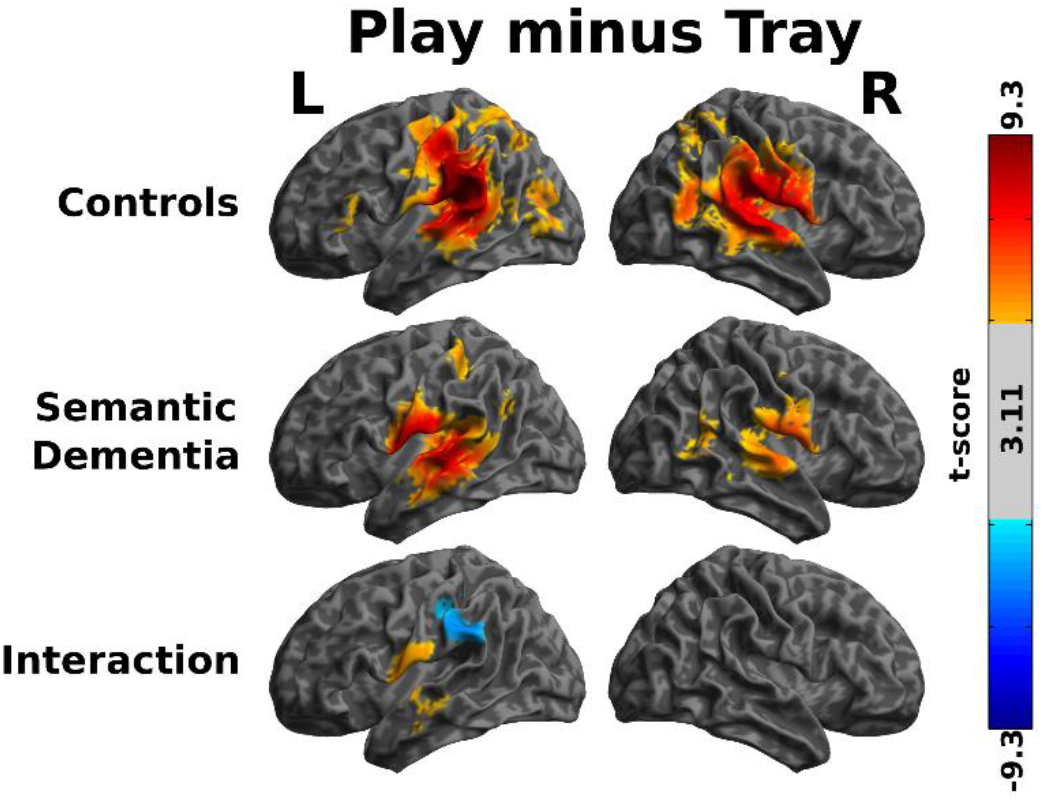
Source reconstructions of the contrast between the two standard real words 50-70ms after onset, the time window during which the largest effect of word identity was demonstrated across the whole brain (cf Figure 6). Shaded areas represent t-scores thresholded at uncorrected p<0.001 (t>3.11). In the row representing the group by condition interaction control responses were subtracted from SD responses such that blue shaded areas represent greater contrast in the controls and red shaded areas represent greater contrast in SD.

**Table 2:**
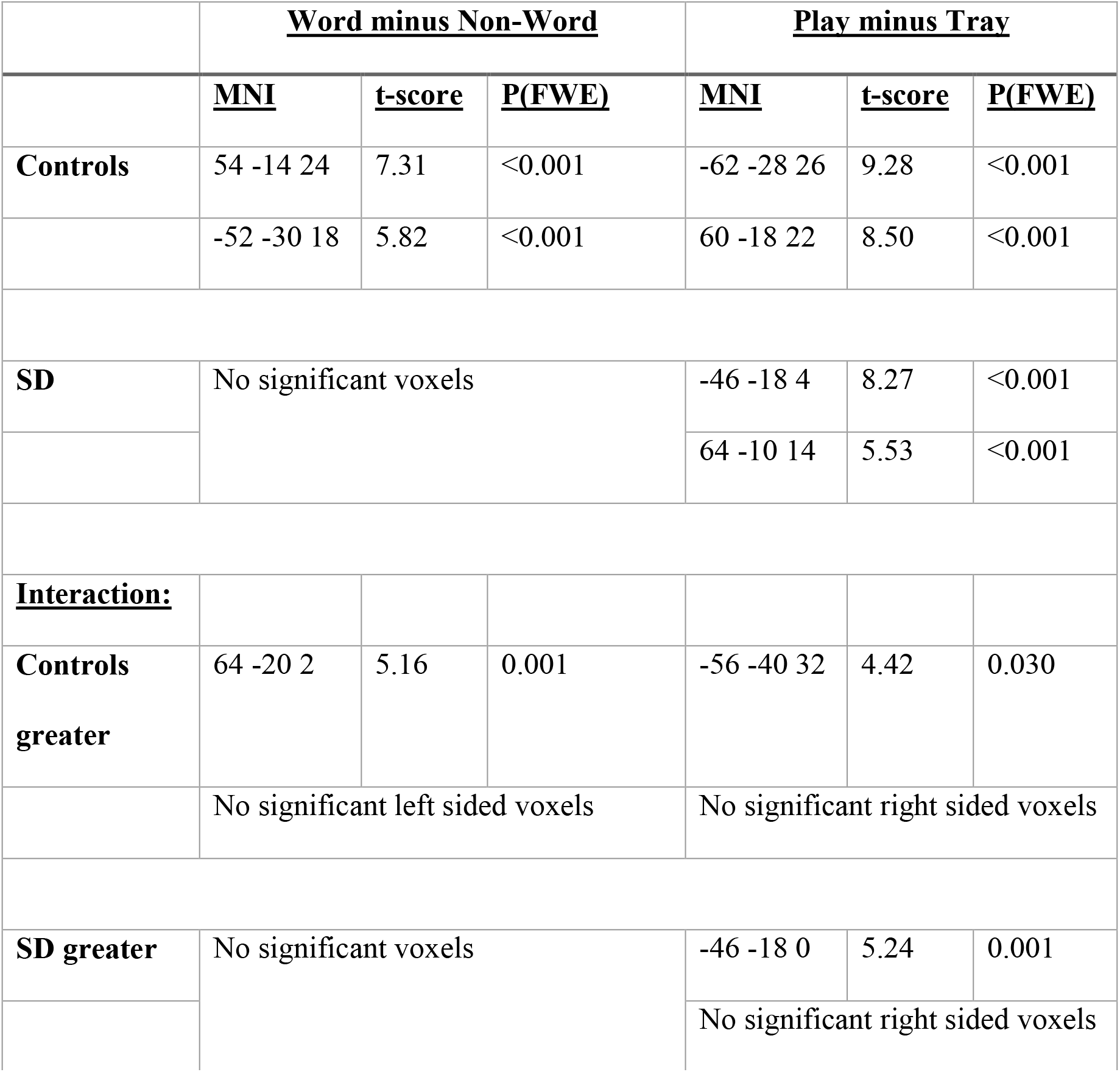
Voxels of peak statistical significance in each hemisphere from the whole-brain corrected SPM contrasts illustrated in Figure 7.

## Discussion

There are three principal results of this study. First, severe degeneration of the left anterior temporal lobe leads to widespread abnormal engagement of right-hemisphere analogues of the language network, during processing of word identity (during 174-294ms, after the divergence point at which stimuli were disambiguated). There was no change in the laterality or magnitude of the peak early response to deviant word endings, occurring approximately 115ms after stimulus disambiguation. This is consistent with a framework in which auditory information passes from primary auditory areas (intact in SD) to ATL so as to engage the left-lateralised processing of word identity. Second, we identified diaschisis – that is, degeneration of the neural architecture in left anterior temporal lobe alters activity in extratemporal brain regions that were not significantly atrophic. Third, we found that in healthy elderly adults, the processing of deviant word endings that change word identity and meaning is strongly left lateralised, as in young healthy adults (Holland *et al*, 2012).

We now relate these results to the three questions posed in the Introduction. First was whether the strongly left-lateralised pattern of activity in healthy controls would shift to a bilateral pattern in the SD patients? This we confirmed. Note that this is not a necessary outcome: of course the brain response in patients will be lower or even largely absent in the lesioned region, but the further consequence of this might be either no increased activity anywhere, or higher responses in other less-damaged left-sided regions. Of particular relevance to the current study, because it was also research in SD, is the fMRI finding by Maguire *et al* (2010) that the usual left-dominant brain activity underlying retrieval of autobiographical memories in controls changed to a pattern of bilateral activity in SD.

A similar question regarding the laterality of brain bases for language processing is often asked (but rarely answered in a definitive manner) in relation to post-stroke aphasia resulting from lesions in classic left-sided language regions—i.e., is it mainly right-hemisphere activity or rather activity in left-hemisphere areas not specialised for language that mediate recovery? The most likely answer is probably that both of these phenomena occur depending on the nature and extent of the lesion (Karbe *et al*, 1998, Price *et al*, 1998). Unsurprisingly, activity in these additional atypical areas does not properly compensate for the reduced response in typical regions: the patients’ performance is always impaired. Although we did not test the SD patients in the current study on their knowledge of the stimulus words, we know from substantial previous research and clinical experience in SD that the patients would easily repeat PLAY or PLAYED or PLATE, but would not necessarily know the words’ identities in the full sense of understanding their meanings. While it seems likely that the patients’ additional right-hemisphere activations contribute to the process of acoustic analysis, preserving word repetition ability, they do not necessarily enable word comprehension.

Our second question was whether we would observe differences in healthy listeners’ brain activity between real, familiar, meaningful words compared to word-like non-words, and whether such differences would be attenuated or absent in SD. Indeed this was the case: our statistical analysis of source-space data (see Figure 7 and Table 2) confirm that where the controls showed significant activation differences in response to words compared to nonwords, the SD patients had no such differences, and there was an interaction between lexical status and diagnostic group. This is in keeping with previous observations suggesting that, in SD, the brain processing of real words and word-like non-words becomes increasingly similar. As mentioned in the Introduction, SD patients are impaired at distinguishing between specially designed words and non-words in visual lexical decision (Rogers *et al*, 2004, Patterson *et al*, 2006). When a real word like FRUIT with rather atypical spelling was paired with a more typically spelled non-word homophone (FRUTE) and the patients were asked to choose the real word, all 22 SD patients had abnormal accuracy, and the more advanced cases tended to prefer the typical non-word to the atypical word as ‘the real thing’. Patterson *et al* (1994) and Knott *et al* (1997) studied immediate serial recall of short word sequences by SD patients, under three conditions: real words that each patient still ‘knew’ or understood; real words that he or she no longer understood; and word-like non-words. Successful recall of the real-but-‘unknown’ words was at a level intermediate between real-“known” words and nonwords. Finally, in tasks of reading aloud briefly presented written words and tasks of identifying words from oral spelling (e.g., “what does C,H,U,R,C,H spell?”), both SD patients and stroke patients with posterior left-hemisphere lesions resulting in pure alexia made many errors (Cumming *et al*, 2006). Strikingly, however, virtually all of the error responses by the pure alexic patients in both tasks were other similar real words, whereas the majority of the errors by the SD patients were orthographically and phonological similar nonwords. All three of these studies were purely behavioural experiments, demonstrating significantly reduced ability to distinguish between real, meaningful words and plausible nonwords. The current study represents an important advance by demonstrating a brain-basis for this phenomenon.

Finally, we asked whether there would be an overall shift in brain activity for the SD patients from areas implicated in normal word processing to those involved in acoustic feature analysis. This too was supported by the findings: group by word interactions demonstrated that the effect of word identity in SD was greater around auditory regions in superior temporal lobe while in controls it was greater around parietal regions (see Figure 8 and Table 2). Although the role of left parietal areas in word processing is not yet well understood or agreed, some authors have suggested that they support the link between auditory processing of words and their meanings (Robson *et al*, 2013, Robson *et al*, 2017), the combination of semantic concepts (Price *et al*, 2015) and the integration of lexical and semantic information (Price *et al*, 2016). It therefore seems likely that, although SD patients have no measurable damage in these caudal and dorsal regions, their significant atrophy in the rostral and ventral temporal lobes would disrupt both forward and backward activations between the two sets of regions, resulting in diaschisis.

In conclusion, therefore, our results indicate that left ATL performs a necessary role in the left-lateralisation of linguistic processing of words, which represents an efficiency saving compared to the bilateral processing of non-words. We suggest that a consequence of ATL atrophy is that automatic word identity processing becomes predominantly acoustic/phonetic rather than lexical.

## Acknowledgements

We thank Lisa Brindley for her assistance in the acquisition of magnetoencephalography data for this study.

The study was primarily funded by the MRC Cognition and Brain Sciences Unit with additional support from the Cambridge NIHR Biomedical Research Centre (the views expressed are those of the authors and not necessarily those of the NHS, the NIHR or the Department of Health and Social Care). TEC was supported by the Association of British Neurologists, and the Patrick Berthoud Charitable trust. YS was supported by the Medical Research Council (MC-A060-5PQ90), Lundbeck Foundation (R164-2013-15801, project 18690), Danish Council for Independent Research (6110-00486, project 23776), and RF P220 framework (No. 14.W03.31.0010). JBR was supported by the Wellcome Trust (103838), and the Medical Research Council (MC-A060-5PQ30/SUAG/004 RG91365).

